# Texans support mountain lion conservation management

**DOI:** 10.1101/2023.08.16.553568

**Authors:** Omar Ohrens, Benjamin Ghasemi, Bonnie E. Gulas-Wroblewski, L. Mark Elbroch

## Abstract

The state of Texas encompasses an estimated 7% of the United States mountain lion (*Puma concolor*) population, a likely overestimate due to their nongame status, unregulated take and high mountain lion mortality rates. In August 2022, Texas Parks and Wildlife Department (TPWD) denied a petition to amend state mountain lion management policies, but was mandated by the Texas Wildlife Commission to form a stakeholder group to assess the potential to initiate mountain lion management and policy. Herein, we synthesize survey data collected and previously published in a report by Ghasemi et al. (2022) to provide members of this mountain lion stakeholder working group, the Commission, and TPWD with succinct summaries of Texas residents’ knowledge and attitudes salient to the evaluation of mountain lion management in the state. We analyzed responses to represent the opinion of all Texans as well as to compare the sentiments of four key stakeholder groups: hunters vs. nonhunters; livestock owners vs. people without livestock; urban vs. rural residents; and Hispanics vs. non-Hispanics. Overall, Texans expressed a positive sentiment about mountain lions and supported management to conserve the species. Respondents exhibited very high consensus regarding the value of scientific research about mountain lions and expressed overwhelming support for mandatory reporting of any mountain lion killed for any purpose by hunters, trappers, or state or federal agents. Texas residents also backed a compensation program supporting livestock producers who lose animals to mountain lions and rapid checking of set traps. Contrary to expectation, hunters and livestock owners were more positive about mountain lions than non-hunters and people without livestock, and we detected no differences in the responses of urban versus rural residents or Hispanics versus non-Hispanics on any topic.

## Introduction

Based upon the International Union for Conservation of Nature (IUCN) Red List of Threatened Species distribution maps (Nielsen et al. 2015), the state of Texas is home to an estimated 7% of the United States’ mountain lion (*Puma concolor*) population. However, Texas likely hosts a smaller proportion due to unregulated take, which has resulted in mountain lion mortality rates that are higher than in other western states (Harveson et al. 2012; Logan and Runge 2021; Elbroch and Harveson 2022), as well as questions about the current persistence of a South Texas mountain lion population (Holbrook et al. 2012; Elbroch and Harveson 2022). In fact, abundance estimates for mountain lion strongholds in west and south Texas are some of the lowest density estimates reported for North America (Harveson et al., 2012). As of 2021, the state wildlife management agency, the Texas Parks and Wildlife Department (TPWD), listed mountain lions as S2/S3 on its Species of Greatest Conservation Need (TPWD 2021), where S2 is an Imperiled classification defined as, “Imperiled in the nation or state/province because of rarity due to very restricted range, very few populations (often 20 or fewer), steep declines, or other factors making it very vulnerable to extirpation from the nation or state/province,” and S3 is a Vulnerable classification defined as “Vulnerable in the nation or state/province due to a restricted range, relatively few populations (often 80 or fewer), recent and widespread declines, other factors making it vulnerable to extirpation.” Despite these worrying trends, mountain lions in Texas continue to be classified as “nongame” and can be killed in any quantity, by any method, and at any time of year. In addition, Texas residents are not required to report any mountain lion harvest or mortality to TPWD.

Chapter 67 of the Texas Parks and Wildlife Code (TPWD 2023b) directs the TPWD to “develop and administer management programs to ensure the continued ability of nongame species of fish and wildlife to perpetuate themselves successfully” (Section 67.002). Further, the Code prescribes that TPWD “conduct ongoing investigations of nongame fish and wildlife to develop information on populations, distribution, habitat needs, limiting factors, and any other biological or ecological data to determine appropriate management and regulatory information” (Section 67.003). Despite inclusion in the Texas list of Species of Greatest Conservation Need, TPWD does not monitor or estimate the abundance of mountain lions within it’s jurisdiction. It could also be argued that TPWD is in violation of Section 34 of the Texas Constitution since the agency has not enacted regulations or other policies to ensure that mountain lion hunting is sustainable for future generations in Texas (Texas Constitution 2023).

In June 2022, a stakeholder working group composed of Texas residents, scientists, advocates, a veterinarian, and other private citizens (https://www.texansformountainlions.org/) submitted a Petition for Rulemaking to TPWD and the Texas Wildlife Commission requesting the following six changes to Texas wildlife policy as governed by TPWD: 1) Conduct a statewide study to identify the abundance, status, and distribution of the mountain lion populations in Texas; 2) Require mandatory reporting of wild mountain lions killed or euthanized for any reason by members of the public, state and federal agents acting in their official capacity, and other wildlife responders; 3) Require trappers employing any form of trap or snare to capture mountain lions to examine their devices at least once every 36 hours, to make mountain lion trapping consistent with current furbearer trapping regulations in Texas; 4) Limit mountain lion take in South Texas to 5 animals per year until TPWD can determine the size and status of the population in this area and a stakeholder advisory group can establish sustainable hunting limits for the region; 5) Prohibit canned hunting of mountain lions, or more specifically, the take of mountain lions that have been restricted from movement during a hunt or prior to a hunt; 6) Form a stakeholder advisory group to collaborate with TPWD to write a mountain lion management plan for Texas.

In August 2022, TPWD denied this Petition but provided a full presentation on the status of mountain lions in the state to the Texas Wildlife Commission. Consequently, the Commission mandated that TPWD fulfill the sixth request in the Petition and form a stakeholder group to assess the merits of the full Petition. In February 2023, TPWD hosted its first mountain lion stakeholder working group meeting, which included parties invited by the agency for reasons that were not disclosed but presumably were related to their identification as representatives for key constituent groups interested and affected by mountain lions and their potential management.

In order to inform the proceedings of subsequent meetings of this stakeholder working group, the Texas Wildlife Commission, and TPWD, we synthesized survey data collected and previously published in a public report (Ghasemi et al. 2022) to generate succinct summaries of the attitudes of Texas residents as relevant to reviewing the Petition and potential management of mountain lions in the state. We also assessed ratios of the percentage of people who strongly supported versus strongly opposed (+2 and -2 on a 5-point Likert scale) trapping-based management, hunting-based management, compensation programs for livestock losses, and mountain lion conservation management more broadly. People with the strongest opinions are those most likely to act on their beliefs (Krosnick & Petty 1995), including actions such as attending and speaking at public meetings, calling a wildlife official or politician, or arguing for a particular viewpoint on social media or in newspapers.

These analyses also provided us with the opportunity to test several hypotheses related to stakeholder perceptions of wildlife management. First, we hypothesized that hunters, livestock owners, and people in rural counties would express less support for mountain lions and mountain lion conservation management than nonhunters, people without livestock, and people who live in urban and suburban counties. Hunters, livestock owners, and rural residents experience higher tangible and intangible “costs” of living with large carnivores and more often face the real and perceived risks posed by mountain lions to livestock and ungulate hunting opportunities (Davenport et al. 2010; Young et al. 2015; Knopff et al. 2016; Elbroch et al. 2017; Mitchell et al. 2018). Secondly, we tested the assumption that the majority of Texans are anti-trapping (Vantassel et al. 2010; White et al. 2021) and, therefore are opposed to trapping-based management and the trapping of mountain lions.

## Methods

### Survey instrument, sampling and analyses

The survey instrument was designed in Qualtrics software and administered online by Qualtrics following a quota system. The goal was to obtain an equal split of respondents drawn from 100% urban/suburban zip codes and the other half drawn from 100% rural zip codes, as defined in the U.S. Census Bureau (2016). Participants from rural areas are generally less abundant than urban participants in online surveys. Data collection was conducted from November 2021 through January 2022, and the survey instrument and procedure were approved by the Texas A&M University Institutional Review Board (No. 2021-1339M).

Participants were required to be at least 18 years of age. They were asked for initial demographic data, including their sex, residence, and ethnicity. Overall, recruited participants represented male:female ratios reported in the 2020 Census for Texas, but not proportional representations of ethnicity among Texans (61% white, 39% Hispanic in Census 2020). Therefore, we weighted the data to match these proportions for our analyses using the ‘SURVEY’ package (Lumley 2021) in R (ver. 3.6.2).

Participants were asked whether they had participated in hunting or trapping of any wildlife in the last 12 months, which we used to define “hunters” versus “nonhunters”. Participants were also asked whether they had participated in any livestock (i.e., cattle, dairy cows, goats, and/or sheep) husbandry in the last 12 months, to define “livestock owners” as distinct from people that did not own or raise livestock. Note that a single person could report any combination in participating as hunter, nonhunter, or livestock producer. For example, someone could be a hunter with livestock or a hunter without livestock.

For each of the questions below, we analyzed all respondents pooled together as our best representation of the attitudes, beliefs, and values of the general public in Texas. Next, we performed the analyses separating four comparative stakeholder ingroups to evaluate any differences in their responses: 1) hunters vs. nonhunters; 2) livestock owners vs. people without livestock; 3) rural vs. suburban/urban respondents; and 4) Hispanics vs. non-Hispanics. Given the large proportion of Texans that are Hispanic or Latino, it is imperative to highlight any potential differences in this stakeholder group’s responses to avoid the historic marginalization of this constituency and to ensure their equitable representation in any forthcoming mountain lion management decisions (Chase et al. 2016).

We used parametric tests in R version 3.6.2 for Mac OS X to relate participant responses (perceptions and Likert scale) with independent variables (e.g., comparative groups – hunters, livestock producers, urban vs rural, ethnicity). T-tests were used when comparing means of two independent groups (e.g., trust in agency between hunters and non-hunters). We used chi-square tests to determine whether there were proportional differences in selecting specific responses among ingroups (e.g., legal status of mountain lions and the four comparative groups). When differences were detected, we employed z-proportions tests to determine which response category differed among comparative groups.

### Knowledge of the current status of mountain lions in Texas

Participants were asked two questions about the current status of mountain lions in Texas. First, they were asked to characterize mountain lion populations as one of five options: 1) Extinct; 2) Endangered; 3) Rare but not endangered; 4) Common; or 5) ‘I don’t know’. Second, participants were asked to select the legal status of mountain lions from among four options: 1) Game animal with controlled seasons regulating when and how many can be hunted or trapped; 2) Non-game animal with no hunting or trapping restrictions; 3) Protected animal that cannot be legally hunted or trapped; or 4) ‘I don’t know’. We analyzed the weighted data to report the proportions of Texas respondents that selected each category, and then used chi-square analyses with a Rao and Scott adjustment (Rao and Scott 1987) to assess for different proportions reported by different ingroups. When differences were detected, we employed z-proportions tests to determine which response category differed among ingroups.

### General sentiment about mountain lions, mountain lion management and research, and the Texas Parks and Wildlife Department

A random sample of participants were asked to respond to statements on the three topics of mountain lions, mountain lion management, and trust in TPWD, resulting in different sample sizes for each analysis. People responded to statements on these topics by choosing from among five options representing a Likert scale ranging from -2 to 2: 1) strongly disagree; 2) somewhat disagree; 3) neither agree nor disagree; 4) somewhat agree; or 5) strongly agree. As described above, we analyzed the weighted data to describe the perspective of the general public in Texas, and we compared the responses among our 4 categories of stakeholder ingroups as well.

For the first topic, we combined the responses of five statements (Table 1) to quantify people’s general sentiments about mountain lions. We used the inverse values for statement 5 so as to combine results with a potential range of -10 to 10. For the second topic, we combined participant responses to four statements (Table 1) to determine whether people in general are supportive of managing mountain lions. We used the inverse values for statement 2 to combine the results of these four statements to create a potential range of -8 to 8. For the third topic, we combined the responses for seven statements that reflect people’s trust in TPWD’s current actions related to mountain lions to create a potential range of -14 to 14. We considered mean responses for these three topics to be unsupportive when their values were negative and their 95% confidence intervals (CIs) did not overlap with 0, neutral if CIs overlapped zero, and to be supportive if their values were positive. We employed t-tests to compare the responses of stakeholder ingroups and considered groups to have responded differently using a 95% significance test (p<0.05).

**Table 1.**
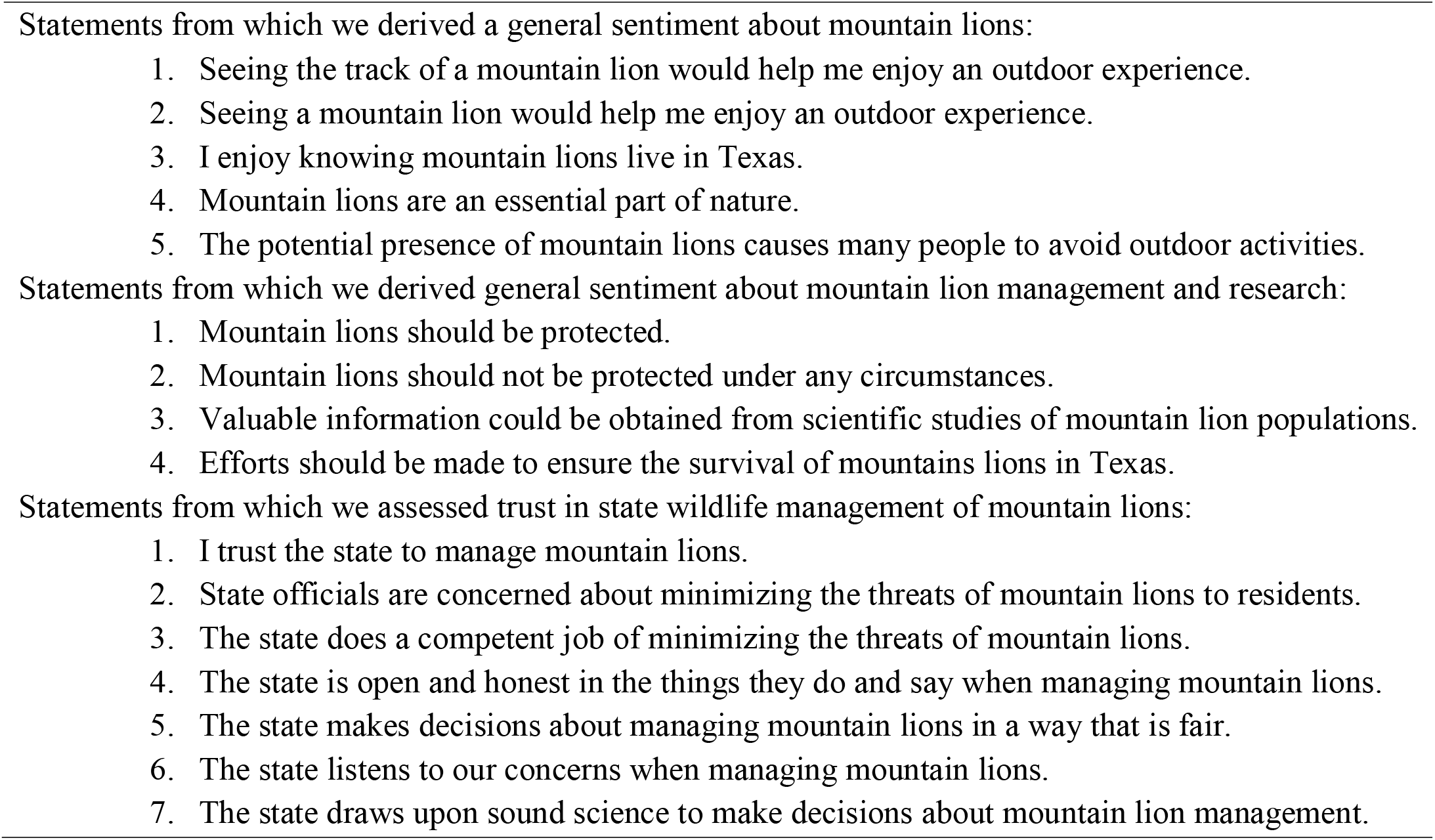
Statements used in our analyses regarding general sentiment about mountain lions, mountain lion management, and trust in the state management of mountain lions.

Following these analyses, we also calculated Potential for Conflict Index (PCI_2_) scores (Vaske et al., 2010) to assess the relative consensus among participants with regards to their responses to individual statements in each of the three topics. PCI_2_ values range from 0 to 1, where 0 indicates complete consensus and zero potential for conflict, and 1 indicates complete disagreement and the highest potential for conflict.

### Trapping regulations and mandatory harvest reporting

Participants were asked to select one of five options related to how often trap regulations should require trappers to check their traps in the field: 1) Daily; 2) 36 hours; 3) 48 hours; 4) 72 hours; or 5) Weekly. Participants were also given the option of three choices to indicate if they thought trap checks should not be required, that trapping should not be allowed, or that they were unsure: 6) There should not be trap checks regulations.; 7) Unsure; or 8) Trapping should not be allowed. In addition, participants were asked whether they support mandatory harvest reporting for mountain lions with three possible responses: 1) yes; 2) no; or 3) unsure. Data were analyzed as described above.

### Other management considerations

We asked three additional questions that provide insight and guidance for decision makers and policy makers with regards to future mountain lion management. We inquired whether 1) Trapping was an acceptable management practice for mountain lions; 2) Hunting was an acceptable management practice for mountain lions, and 3) Whether the participant would endorse a management plan inclusive of a compensation program for livestock producers that lose animals to mountain lions. All responses were recorded on a 5-point Likert scale (range from -2 or strongly disagree, to + 2 or strongly agree). Responses were analyzed individually to describe the general response of all people surveyed, then followed with analyses to assess any differences detected among our four ingroups of interest. Analyses were conducted per descriptions above.

## Results

### Participants

Qualtrics secured 740 respondents but were unable to meet the quotas set by the researchers responsible for the survey: 237 people reported living in rural zip codes (Figure 1), 89 reported having participated in hunting or trapping in the previous 12 months, 87 reported owning or raising livestock in the previous 12 months, and 141 self-identified as Hispanic. We weighted the data to account for an underrepresentation of Hispanic people among the respondents, which reduced the sample size to 702 people in terms of analytics.

**Figure 1.**
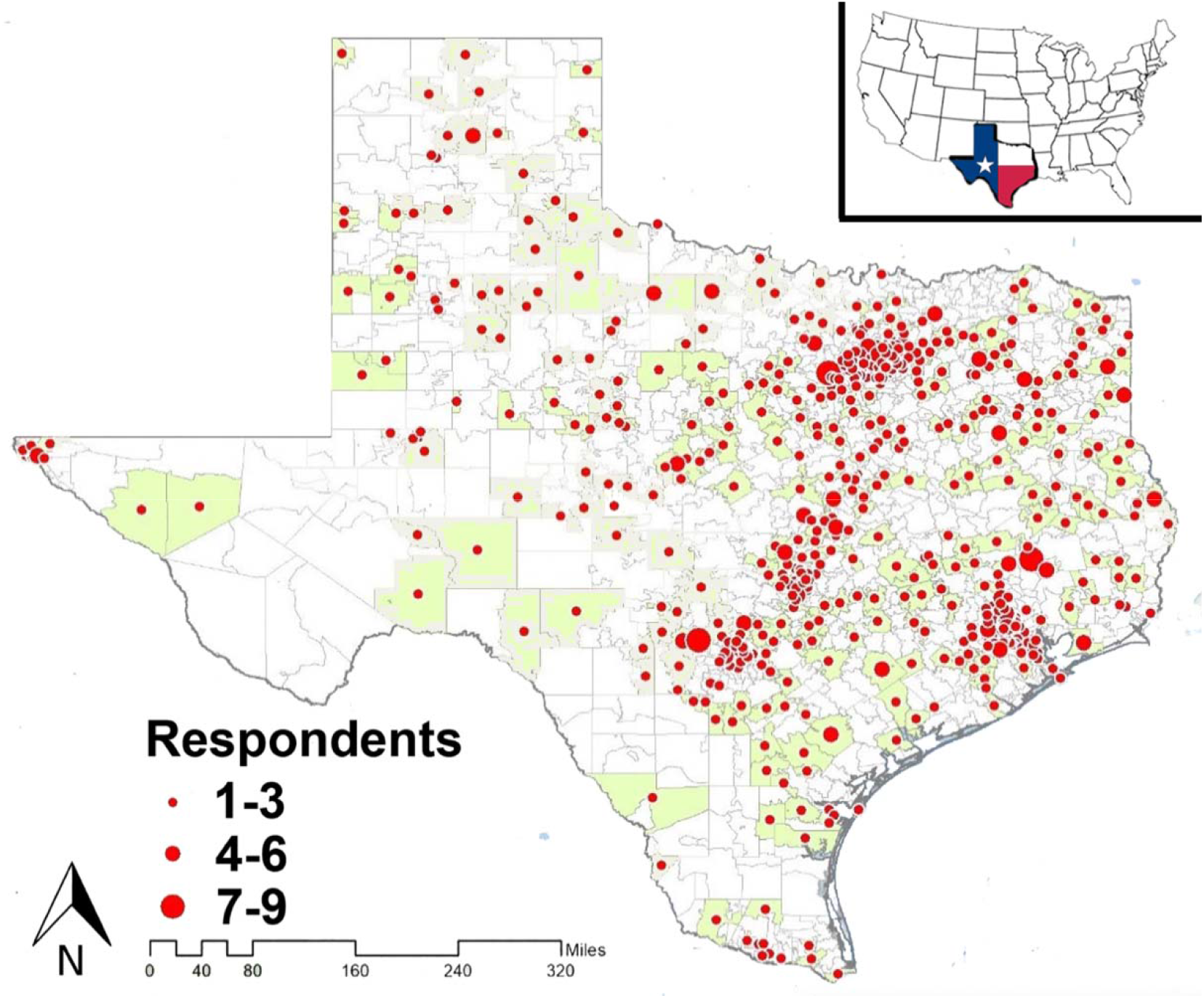
Inset shows the location of Texas in the contiguous United States. Main map displays the geographical distribution of survey respondents within Texas (Ghasemi et al. 2022).

### Knowledge of current status of mountain lions in Texas

More people in Texas correctly reported that mountain lions were rare but not endangered (32%) than any other response, and 28% believed mountain lions were endangered. Twenty-one percent reported that they did not know the status of mountain lions, 13% reported that they were common, and 5% indicated that they were extinct. There were differences in the proportional responses for hunters versus non-hunters (F_3.89, 2725.0_ = 2.830, p = 0.025). More hunters incorrectly responded that mountain lions were common than non-hunters (hunter = 23%, nonhunter = 12%, *z* = 2.826, p = 0.005), and more non-hunters reported that they did not know the current status of mountain lions in Texas (hunter = 11%, nonhunter = 23%, *z* = -2.312, p = 0.009) (Table S1). There were also differences in the proportional responses of livestock owners versus people without livestock (F_4.0, 2787.2_ = 4.422, p = 0.001). More livestock owners incorrectly reported that mountain lions were common than people without livestock (livestock = 20%, no livestock = 12%, *z* = 1.979, p = 0.048), and more livestock owners also incorrectly reported that mountain lions were extinct (livestock = 13%, no livestock = 5%, *z* = 3.075, p = 0.002) (Table S2). More people without livestock reported that they did not know the current status of mountain lions in Texas (livestock = 9%, no livestock = 23%, *z* = -2.905, p = 0.003). There were no differences in proportional responses among rural versus urban inhabitants (F_4.0, 2799.2_ = 0.577, p = 0.679) nor between Hispanics versus non-Hispanics (F_4.0, 2801.1_ = 1.527, p = 0.192).

**Table S1.**
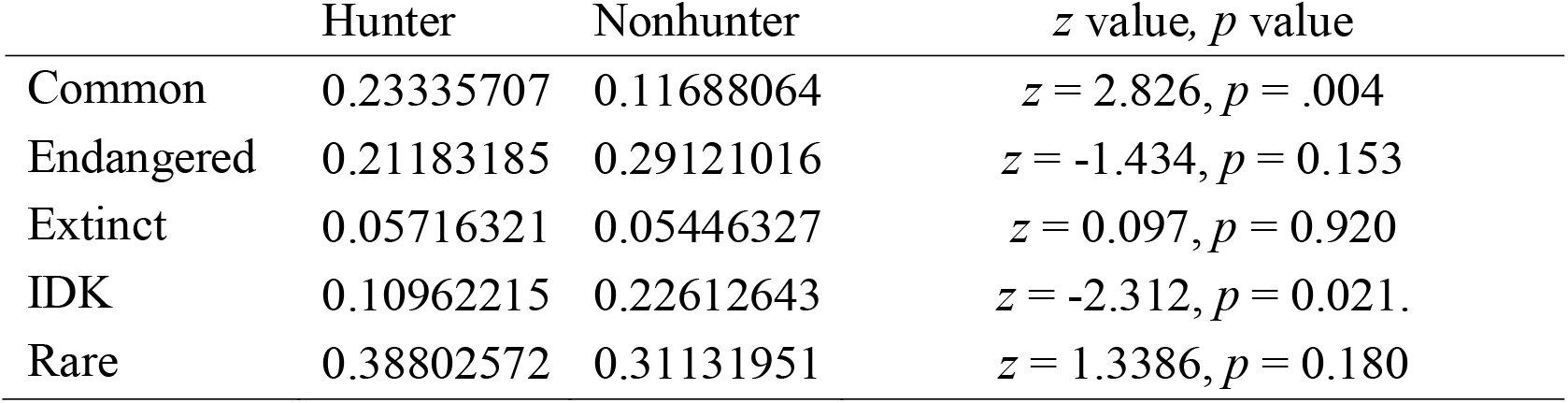
Proportional responses of hunters and nonhunters with regards to the current population status of mountain lions in Texas with accompanying *z* and *p* values. IDK = I don’t know.

| | Hunter | Nonhunter | $z$ value, $p$ value |
| --- | --- | --- | --- |
| Common | 0.23335707 | 0.11688064 | $z = 2.826$, $p = .004$ |
| Endangered | 0.21183185 | 0.29121016 | $z = -1.434$, $p = 0.153$ |
| Extinct | 0.05716321 | 0.05446327 | $z = 0.097$, $p = 0.920$ |
| IDK | 0.10962215 | 0.22612643 | $z = -2.312$, $p = 0.021$. |
| Rare | 0.38802572 | 0.31131951 | $z = 1.3386$, $p = 0.180$ |

**Table S2.** Proportional responses of livestock owners and people without livestock with regards to the current population status of mountain lions in Texas with accompanying *z* and *p* values. IDK = I don’t know.

| | Livestock | No livestock | $z$ value, $p$ value |
| --- | --- | --- | --- |
| Common | 0.19845379 | 0.12006916 | $z = 1.979, p = 0.048$ |
| Endangered | 0.2121016 | 0.2921237 | $z = -1.5039, p = 0.134$ |
| Extinct | 0.12775326 | 0.04516623 | $z = 3.075, p = 0.002$ |
| IDK | 0.08944461 | 0.23016745 | $z = -2.905, p = 0.004$ |
| Rare | 0.37224674 | 0.31247345 | $z = 1.085, p = 0.276$ |

Overall, the majority of people in Texas (45%) incorrectly believed mountain lions are a protected species that cannot be hunted or trapped. There were, however, differences in the proportional responses for hunters versus non-hunters (F_3.00, 2100.7_ = 5.496, p = 0.001). More hunters than non-hunters incorrectly believed that mountain lions were a game species in Texas (hunter = 18%, nonhunter = 9%, *z* = 2.440, p = 0.015), and, at the same time, more hunters than non-hunters correctly believed that mountain lions were a nongame species with unmanaged harvest or reporting (hunter = 32%, nonhunter = 17%, *z* = 3.050, p = 0.002) (Table S3). In contrast, a larger proportion of nonhunters incorrectly believed that mountain lions are a protected species (hunter = 29%, nonhunter = 48%, *z* = -2.983, p = 0.003).There were no differences in proportional responses among livestock owners and people without livestock (F_2.99, 2093.8_ = 2.143, p = 0.093), rural versus urban inhabitants (F_3.0, 2100.2_ = 0.125, p = 0.945), or Hispanics versus non-Hispanics (F_3.0, 2101.7_ = 1.666, p = 0.172).

**Table S3.** Proportional responses of hunters and nonhunters with regards to the current management status of mountain lions in Texas with accompanying *z* and *p* values. IDK = I don’t know.

| | Hunter | Nonhunter | $z$ value, $p$ value |
| --- | --- | --- | --- |
| Game | 17.8% | 8.8% | $z = 2.440, p = 0.015$ |
| IDK | 20.9% | 26.2% | $z = -0.981, p = 0.327$ |
| Nongame | 31.9% | 17.2% | $z = 3.050, p = 0.002$ |
| Protected | 29.4% | 47.7% | $z = -2.983, p = 0.003$ |

### General sentiment about mountain lions

People in Texas exhibited a slightly more positive than neutral sentiment about mountain lions (mean score = 0.38 *±* 0.08, scale -10 to +10). Hunters expressed a more positive sentiment about mountain lions (mean score = 0.98 ± 0.30 SE) than nonhunters (mean score = 0.31 *±* 0.08 SE) (t_700_ = -2.116, p = 0.03). Similarly, livestock owners exhibited a more positive sentiment about mountain lions (mean score = 0.93 ± 0.28 SE) than those who do not own livestock (mean score = 0.31 *±* 0.08 SE) (t_700_ = -2.104, p = 0.04). There were no significant differences in sentiment among rural versus urban respondents (t_700_ = 0.942, p = 0.347) or Hispanic versus non-Hispanic respondents (t_700_ = 0.138, p = 0.889).

The ratio of strongly agree:strongly disagree for the statement describing mountain lions as an essential part of nature was 35:4, but ratios were more in favor of strongly disagree with regards to feeling excited about detecting evidence of mountain lions in the field (12:17 for finding footprints, 16:17 for seeing the animal) (Table S4). This pattern was also clear in terms of general consensus among participants. Texans exhibited the highest consensus in their belief that mountain lions are an essential part of nature (PCI = 0.16) and the least consensus (i.e., the greatest potential for conflict) regarding statements about the influence of seeing the footprint of a mountain lion (PCI = 0.31) or the actual animal (PCI = 0.33) on one’s outdoor experience (Figure 2).

**Table S4.** Percentages of people who responded for each category for each of 5 statements reflecting general sentiment about mountain lions.

|  | Strongly disagree | Somewhat disagree | Neutral | Somewhat agree | Strongly Agree |
| --- | --- | --- | --- | --- | --- |
| Seeing track of mountain lion would help enjoy outdoor experience | 17% | 16% | 30% | 25% | 12% |
| Seeing a mountain lion would help enjoy outdoor experience | 17% | 13% | 28% | 27% | 16% |
| Enjoy knowing mountain lions live in Texas | 10% | 8% | 34% | 26% | 23% |
| Mountain lions are essential part of nature | 4% | 4% | 15% | 41% | 35% |
| Potential presence of mountain lions causes people to avoid outdoor activities | 8% | 15% | 29% | 33% | 15% |

**Figure 2.**
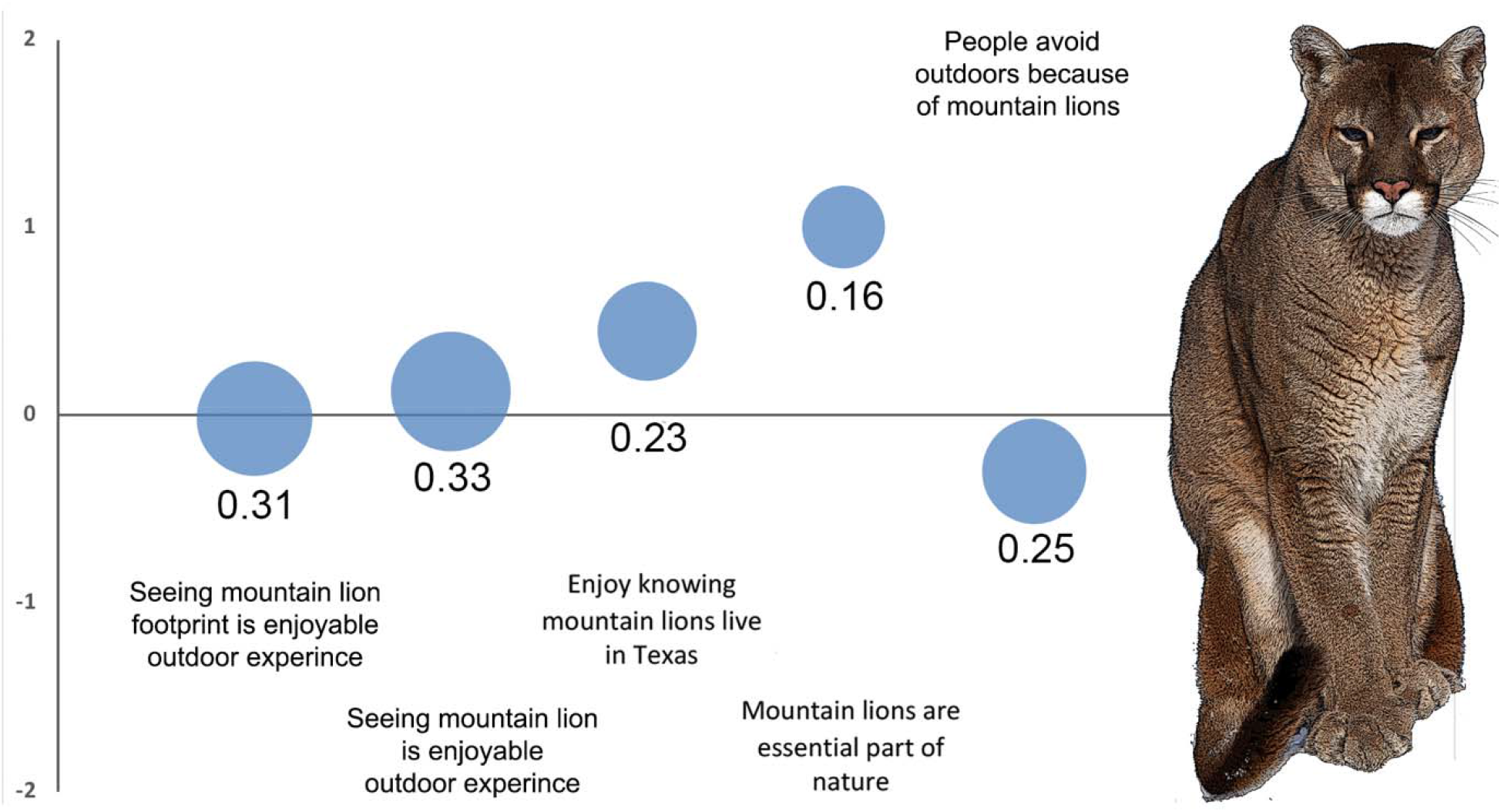
Mean scores for each of five statements reflecting general sentiment about mountain lions and their related PCI scores. The larger the PCI value and the size of the circle, the less consensus among respondents and the greater potential for conflict. The focal species, *Puma concolor*, is pictured on the right.

### Sentiment about mountain lion management

The people of Texas expressed slightly positive support for managing mountain lions (mean score = 1.04 *±* 0.06, scale = -8 to 8). Based on our p<0.05 threshold, there were no differences in responses among hunters and nonhunters, although the results were marginal (hunter score = 0.69 *±* 0.19, nonhunter score = 1.08 *±* 0.06, t_700_ = 1.881, p = 0.06). We also did not find differences between livestock owners and people without livestock (t_700_ = 0.118, p = 0.905), rural versus urban inhabitants (t_700_ = -0.466, p = 0.641), or Hispanic versus non-Hispanic respondents (t_700_ = 0.834, p = 0.404).

Overall, there were a much larger proportion of participants who “strongly agreed” with statements expressing positive sentiment about mountain lion management than there were those who “strongly disagree” with mountain lion management (Table S5). For example, for every 32 people who strongly agreed with the statement that mountain lions should be protected, there were five people that strongly disagreed with the statement. Similarly, for every 31 people who strongly agreed with the statement that efforts should be made to ensure the species’ survival in Texas, five people strongly disagreed with this assertion.

**Table S5.**
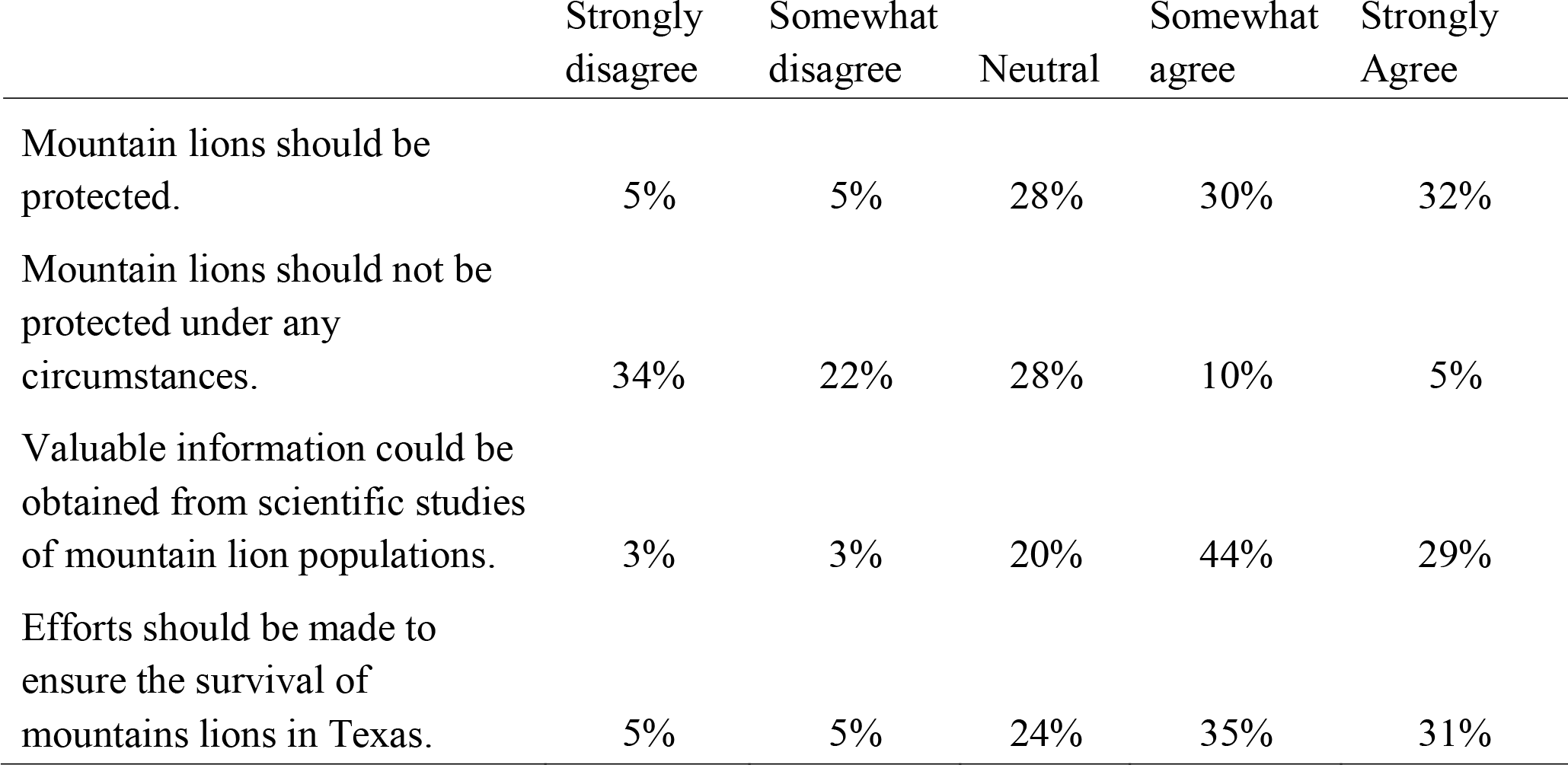
Percentages of people who responded for each category for each of four statements reflecting general sentiment about mountain lion management.

In relation to mountain lion management, Texans exhibited the highest consensus (PCI = 0.11) in their belief that science and research are important. Texas respondents demonstrated the least consensus around the statement that mountain lions should not be protected under any circumstance (PCI = 0.23) (Figure 3).

**Figure 3.**
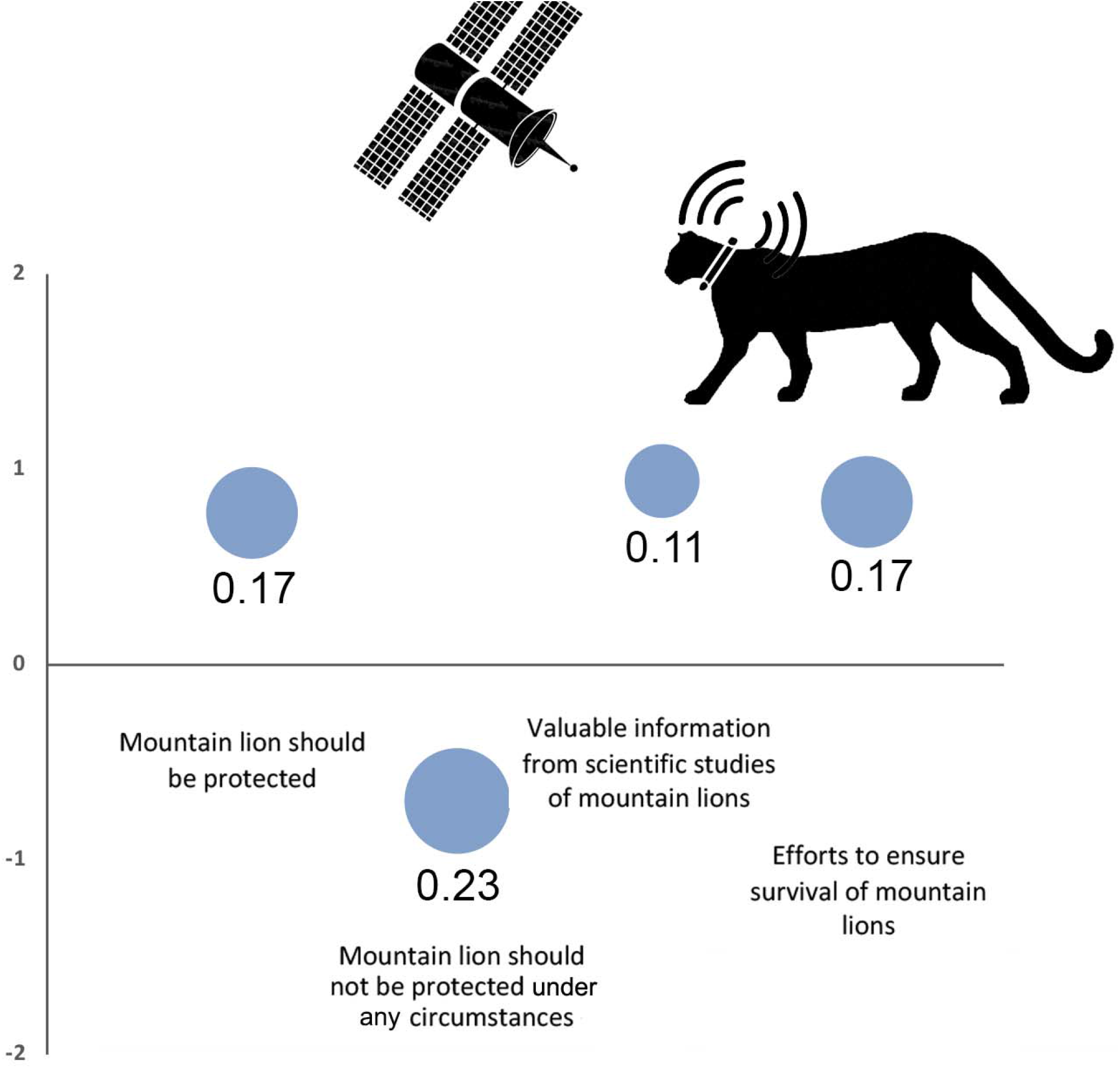
Mean scores for each of four statements reflecting general sentiment about mountain lion management and their related PCI scores. The larger the PCI value and the size of the circle, the less consensus among respondents and the greater potential for conflict.

### Assessing trust in the Texas Parks and Wildlife Department to effectively manage mountain lions

In general, Texans exhibited modest trust in TPWD in relation to managing mountain lions and employing the best available science to guide decisions about mountain lion management (mean score = 2.23 *±* 0.23 SE, scale = -14 to +14). However, hunters (score = 3.47 *±* 0.62 SE) expressed greater positive sentiment about TPWD than did nonhunters (score = 2.08 *±* 0.24 SE) (t_700_ = -2.082, p = 0.03). Livestock owners (score = 3.46 *±* 0.63 SE) also exhibited greater positive sentiment for TPWD than did people without livestock (score = 2.07 *±* 0.24) (t_700_ = -2.075, p = 0.03). There were no significant differences in trust at the p<0.05 threshold among rural versus urban inhabitants (t_700_= -0.746, p = 0.455) nor among Hispanics versus non-Hispanics (t_700_= 1.736, p = 0.08).

Overall, Texas residents demonstrated medium to high consensus in their trust towards TPWD (Figure 4). In particular, there was high consensus for the statement that TPWD makes scientifically sound decisions about managing mountain lions (PCI = 0.09). Conversely, there was less consensus (PCI = 0.15) and a more neutral view expressed related to the statement about how honest and transparent TPWD is in their management of mountain lions (Figure 4).

**Figure 4.**
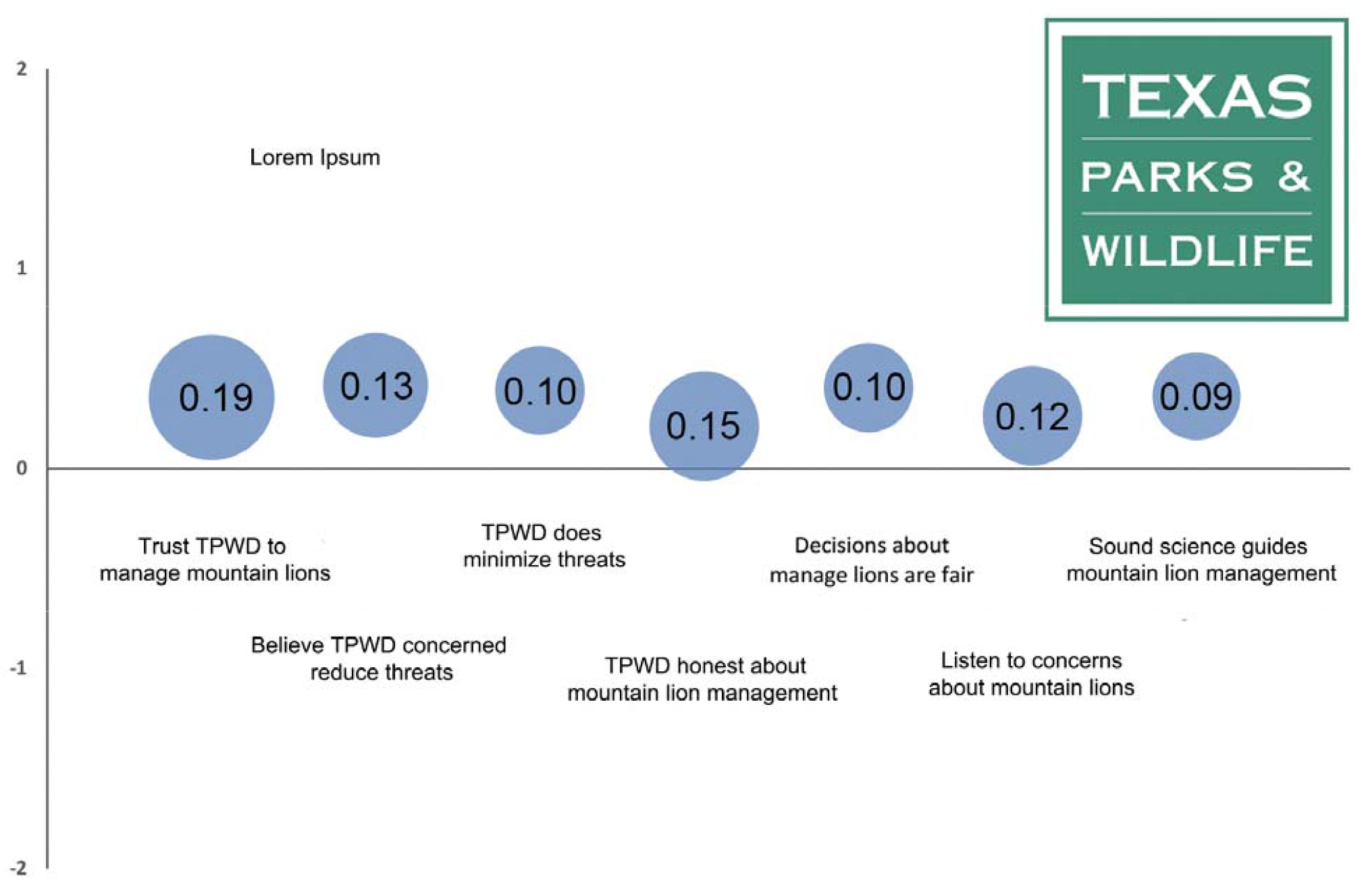
Mean scores for each of seven statements reflecting general sentiment about the trustworthiness of TPWD and their related PCI scores. The larger the PCI value and the larger the circle, the less consensus among respondents and the greater potential for conflict.

### Trapping regulations and mandatory harvest reporting

The people of Texas overwhelmingly supported daily trap checks for mountain lions. When indicating their preferred rate of trap checks, 64% of respondents selected daily trap checks, 11% selected every 36 hours (the current trapping regulation for furbearers in Texas), 6% selected every 48 hours, 1% selected every 72 hours, 6% were unsure, 1% expressed that there should not be trap checks, and 7% expressed that trapping should not be allowed. There were no significant differences in the proportional selection in the eight answer categories among hunters versus nonhunters (F_6.3, 4399.6_ = 1.169, p = 0.319), livestock owners vs people without livestock (F_6.6, 4638.0_ = 0.672, p = 0.687), rural versus urban inhabitants (F_6.9, 4856.5_ = 0.848, p = 0.546), or Hispanics versus non-Hispanics (F_6.9, 4869.0_ = 0.974, p = 0.448).

Texas residents strongly approved the reporting of any mountain lions killed or euthanized in the state. Seventy-four percent of respondents supported harvest reporting, 18% were unsure, and 8% did not support harvest reporting. There were differences in the proportional responses for hunters versus nonhunters (F_1.89, 1327.0_ = 5.423, p = 0.005). Hunters and nonhunters expressed equal support for harvest reporting (hunter = 77%, nonhunter = 74%, *z* = 0.648, p = 0.516), but more hunters opposed harvest reporting than did nonhunters (hunter = 15%, nonhunter = 7%, *z* = 2.597, p = 0.009), and less hunters were unsure than were nonhunters (hunter = 7%, nonhunter = 19%, *z* = -2.534, p = 0.011). There were no significant differences in proportional selection in the three answer categories among livestock owners versus people without livestock (F_2.0, 1396.6_ = 0.443, p = 0.641), rural versus urban inhabitants (F_2.0, 1401.3_ = 0.251, p = 0.778), or Hispanics versus non-Hispanics (F_2.0, 1401.9_ = 0.936, p = 0.393).

### Other management considerations

Respondents were neutral (mean score = 0.057 *±* 0.09 SE) with regards to the use of trapping as a management tool for mountain lions in Texas. There were no differences of opinion among any stakeholder ingroup that was tested. For every seven people that are strongly opposed to trapping as a management tool, there were six people who indicated strong support for this type of management (Table S6).

In comparison, Texas residents (mean score = -0.103 *±* 0.09 SE) were just slightly negative of neutral with regards to the use of hunting as a management tool for mountain lions. Hunters (mean score = 0.83 *±* 0.20) expressed support for hunting as a management tool, whereas nonhunters (mean score = -0.22 ± 0.09) (t231 = -4.843, p <0.001) were slightly against hunting as a management option. Livestock producers (mean score = 0.30 *±* 0.21) similarly expressed a slightly positive view of hunting as a management tool, whereas people without livestock (mean score = -0.16 *±* 0.09) (t_231_ = -1.978, p = 0.049) were slightly opposed to management via hunting. There were no significant differences of opinion between rural versus urban residents (t_231_ = 0.752, p = 0.452) or Hispanics and non-Hispanics (t_231_ = -0.566, p = 0.572).

Texans expressed support for a compensation program to aid livestock owners that lose animals to mountain lions (mean score = 0.570 ± 0.07). For every person who strongly opposed such a program, four expressed strong support for livestock compensation initiatives (Table S6). There were no significant differences among responses made by our four ingroups of interest (hunter versus nonhunter, t = -1.489, p = 0.138; livestock versus no livestock, t = -0.756 p = 0.450; rural versus urban, t = -0.450, p = 0.653; and non-Hispanic versus Hispanic, t = 0.076, p = 0.94).

**Table S6.** Proportional responses (%) for each of three statements about potential mountain lion management tools.

|  | Strongly disagree | Somewhat disagree | Neutral | Somewhat agree | Strongly Agree |
| --- | --- | --- | --- | --- | --- |
| Trapping is an acceptable way of managing mountain lion populations. | 14 | 12 | 36 | 25 | 12 |
| Hunting is an acceptable way of managing mountain lion populations. | 18 | 18 | 32 | 21 | 11 |
| I would endorse a state management plan that compensates livestock producers for livestock loss from mountain lion predation. | 5 | 6 | 35 | 34 | 20 |

## Discussion

Overall, survey respondents expressed a positive, albeit modest, sentiment towards mountain lions and supported state management of the species. The Texas populace demonstrated a very high consensus in their valuation of scientific research about mountain lions and the need for science to guide management decisions by TPWD. They also overwhelmingly expressed support for mandatory reporting of any mountain lion killed for any purpose by hunters, trappers, or state or federal agents, and for daily checks of traps set in the field. An overwhelming majority endorsed the implementation of a livestock compensation program for producers who lose animals to mountain lions.

These analyses provide definitive, scientific evidence that the people of Texas, inclusive of hunters and livestock producers, support the first three proposed activities in the 2022 Petition for Rulemaking received by the TPWD: 1) Conduct a statewide study to identify the abundance, status, and distribution of the mountain lion populations in Texas; 2) Require mandatory reporting of wild mountain lions killed or euthanized for any reason by members of the public, state and federal agents acting in their official capacity, and other wildlife responders; 3) Require trappers employing any form of trap or snare to capture mountain lions to examine their devices at least once every 36 hours, to make mountain lion trapping consistent with current furbearer trapping regulations in Texas. Likewise, it can be deduced from these survey results that there would most likely be endorsement of the fourth proposed activity that advocates for the use of the best available science to manage Texas’ mountain lion population: 4) Limit mountain lion take in South Texas to 5 animals per year until TPWD can determine the size and status of the population in this area. Nevertheless, this survey did not directly assess the attitudes of the Texas citizenry in relation to either activities Four or Five of the 2022 Petition.

Hunters versus nonhunters and livestock producers versus people without livestock exhibited some differences in support for and sentiment about mountain lions and mountain lion management. However, these discrepancies were not entirely in alignment with our initial predictions based on the existing literature. Hunters and livestock owners expressed greater trust in state wildlife management, a trend that we hypothesized given that these stakeholders are generally more engaged with state wildlife agencies and predominantly viewed as the primary constituents of state wildlife programs (Adams et al. 1997; Decker et al. 2019). Contrary to our expectations, hunters and livestock producers expressed more positive sentiments for mountain lions and their management than did nonhunters and people without livestock. These attitudes are noteworthy, considering that state agencies often believe they are acting on behalf of their constituents when they reduce large carnivore populations or implement more liberal harvest management of these species (e.g., Mitchell et al. 2018; Profitt et al. 2020).

Furthermore, we detected no significant differences in the responses between rural and urban respondents, dispelling the assumption that there are clear attitudinal divisions between people living sympatric with large carnivores in rural areas and those living more isolated from large carnivores in urban or suburban environments (Williams et al. 2002). We also detected no differences between the overall responses of Hispanic and non-Hispanic participants, and we did not find strong opposition to trapping-based management among Texans in general. The later result was likely due to the broad framing of the question, which enabled “trapping management” to include diverse trapping activities, ranging from controlling invasive species to protect local resources to recreational trapping for fur sales to international markets. In contrast, survey respondents exhibited slightly negative sentiment for hunting-based management, an unexpected result considering that our participant pool included a high representation of hunters (12% of participants in this survey were hunters in comparison to a previous report that only 4% of Texans hunt; Stacker 2021).

In conclusion, Texans expressed positive sentiments for mountain lions, the management of this species in Texas, and their trust of TPWD, though their responses were only slightly better than neutral. In part, these attitudes may be attributable to the perceived or real risks of coexisting with mountain lions, such as the fear of mountain lions reflected in the proportional negative versus positive responses about finding mountain lion tracks in the field or seeing the actual animal (Table S4). This may further be indicative of a general disengagement with the state wildlife agency, wildlife topics in general, or perhaps in relation to mountain lions specifically, an inference substantiated by the fact that the majority of respondents misclassified the current management and population status of mountain lions in Texas. The majority of people also thought that mountain lions were a protected species in Texas. We believe TPWD must prioritize public outreach and education to a greater diversity of Texans to 1) mitigate the perceptions of risks and/or costs associated with mountain lions (*sensu* Johansson et al. 2017), 2) increase public knowledge about the species and its management, and 3) simultaneously foster greater trust in natural resource governance in order to prevent disengagement by the public (Rapp 2020). Furthermore, we strongly encourage the mountain lion stakeholder working group, the Texas Wildlife Commission, and TPWD to incorporate these findings into their evaluation of the 2022 Petition and potential mountain lion policies to ensure that the state’s mountain lion management decision-making is based on the best available science and equitably represents the diverse perspectives of all Texas constituents to create a genuinely adaptive and integrated management plan to preserve the future of mountain lions, Texas communities, and the ecosystems on which we all depend.

## Acknowledgements

The original survey was funded by a grant from the Summerlee Foundation (grant #M2200612). We would also like to express our gratitude to Monica Morrison and Karin Saucedo for their insights in the early stages of this project.

